# PMA-qPCR: ACCELERATING THE MARKET RELEASE OF HIGH-QUALITY *BRADYRHIZOBIUM DIAZOEFFICIENS* INOCULANTS

**DOI:** 10.1101/2025.05.22.655534

**Authors:** Mariana Cap, Camila Frydman, Antonella Galiñanes, Camila Aranguiz, Irina Faraco, Luisina Andriolo, Viviana Parreño, Marina Mozgovoj

## Abstract

The quantification of *Bradyrhizobium diazoefficiens* in inoculants traditionally relies on culture-based assays, which are labor-intensive and time-consuming. To address this limitation, we validated a PMA-qPCR assay as a rapid and reliable alternative for estimating viable *Bradyrhizobium* counts. The assay demonstrated strong performance, achieving approximately 95% efficiency, a standard deviation of 0.3 log CFU/ml, and an intra-assay reproducibility with a coefficient of variation less than 10%. Key experiments optimized PMA concentration to ensure selective exclusion of non-viable cells without compromising viable cell quantification. Discrimination threshold assessment confirmed the assay’s ability to differentiate quarter-strength dilutions. Final validation against plate counting revealed an 82% correlation, significantly reducing processing time from 120 hours to just 5 hours. This is the first study to apply PMA-qPCR specifically for *Bradyrhizobium diazoefficiens* quantification in inoculants. The results highlight its potential as a high-throughput tool for microbial viability assessment, offering improvements in efficiency and precision for batch-to-batch quality control in industrial applications.

## 1. Introduction

The increasing concern over climate change and the need for more sustainable agricultural practices have positioned *Bradyrhizobium* as a valuable tool for legume inoculation. These bacteria establish symbiotic relationships with plants by forming root or stem nodules, where they fix atmospheric nitrogen and convert it into ammonia, a form readily available for plant uptake. However, the growing demand for *Bradyrhizobium* inoculants has revealed a major bottleneck in the production process: the lengthy and labor-intensive viability certification [1,2]. Currently, the viability assessment relies on plate counting, a method that requires several days of incubation, delaying the product’s release to the market. This creates significant pressure on inoculant manufacturers to optimize their processes and adopt more efficient certification methods. To overcome the challenge of ensuring both rapid product delivery and compliance with strict quality standards, the use of PMA-qPCR was evaluated.

PMA stands for Propidium Monoazide which is a membrane-impermeant dye that binds to DNA in non-viable cells. The basic mechanism relies on the ability of PMA to penetrate compromised membranes of non-viable cells, given that when the cell membrane is intact, PMA is unable to penetrate. A following photoactivation step converts the azido group of the PMA to a reactive nitrene radical which reacts with the double-stranded DNA and binds to it with high affinity [3]. This reaction inhibits PCR amplification of the modified DNA-targeted sequence [4]. This approach has been successfully applied by other researchers for various matrices and bacterial species. For instance, Bouju-Albert et al. (2021), validated PMA-qPCR for quantifying viable *Brochotrix thermosphacta* in cold-smoked salmon. Shi et al. (2022), used PMA-qPCR to selectively quantify viable lactic acid bacteria in fermented milk. In this regard, we demonstrated the effectiveness of PMA-qPCR in detecting and quantifying viable Shiga toxin-producing *Escherichia coli* in beef burgers after treatment with high hydrostatic pressure [7,8].

This study aimed to evaluate the effectiveness of PMA-qPCR for quantifying viable *Bradyrhizobium diazoefficiens* in commercial inoculants. To achieve this, we first optimized the qPCR assay for bacterial quantification, then refined the PMA-qPCR protocol, and finally validated the methodology using the final product.

## 2. Materials and methods

### 2.1 Bacterial strain, growth conditions and traditional microbiological quantification

*Bradyrhizobium diazoefficiens (strain SEMIA 5080)*, provided by Rizobacter S.A. culture collection, was used in this study for the assay standardization. For quantification using the plate count method, bacterial cultures were serially diluted, and spread onto modified Mannitol Yeast Agar with Congo Red (MYA+CR). Plates were incubated at 29.5±1°C for 4-5 days or until colonies reached a diameter of 1.5 to 3 mm. Results were expressed as the Log_10_ of colony-forming units (CFU) per ml.

### 2.2 DNA Extraction

Genomic DNA was extracted using Tian amp Bacteria DNA kit (Tiangen, China) following the manufacture instructions for Gram negative bacterial DNA.

### 2.3 qPCR

Primers and TaqMan probe were designed based on conserved regions identified through sequence alignment of *Bradyrhizobium diazoefficiens* strains SEMIA 5080 (accession NZ_ADOU02000008.1) using the Nucleotide BLAST tool. Primers and probe sequences were as follows: primer FW 5’GGAAGCAAGCGACAAGTCT 3’; primer REV 5’TTCGTGCTCGACAATCTCAC 3’ and Probe 5’FAM-TCGTTAACCATCCGTCAACCAGCA-MGB-EDQ 3’. Real-time PCR was performed using the StepOnePlus™ System (Thermo Fisher Scientific, USA). The reactions volume was 12.5 µl volume, containing 5 µM of 2X iTaq Universal Probes Supermix (Bio-Rad, USA); 0.5 µM of each primer, 0.25 µM of the probe, and 4.7 µl of DNA template. The thermal cycling parameters were 95°C for 5 min followed by 40 cycles at 95°C for 15 s and 60°C for 1 s.

### 2.4 Optimization of qPCR for Bradyrhizobium diazoefficiens quantification

Initially, a total of six independent assays were conducted using ten serial 5-fold dilutions of *Bradyrhizobium diazoefficiens* DNA, spanning a range from 8.74 to 2.45 log CFU/ml. Each assay was performed in triplicate, resulting in 18 replicates per concentration and a total of 180 measurements. These results were used to estimate the qPCR limit of detection (LOD), limit of quantification (LOQ), dynamic range, and amplification efficiency were estimated following the MIQE guidelines [9,10]. For repeatability and reproducibility, all quantification curves used in the assays were included. To assess the agreement between qPCR-based estimates and traditional plate counts, a simple linear regression model was applied. The LOD was calculated by analyzing the results of the 10 runs of the standard curve, as described in Figure 1a. The value of 1 CFU/ml was included as the baseline of the dose–response curve and was considered non-detectable. Samples were considered positive when Ct values were below 40, based on the upper limit of the prediction interval established for the relationship between Ct value and bacterial plate counts (CFU/ml).

**Figure 1.**
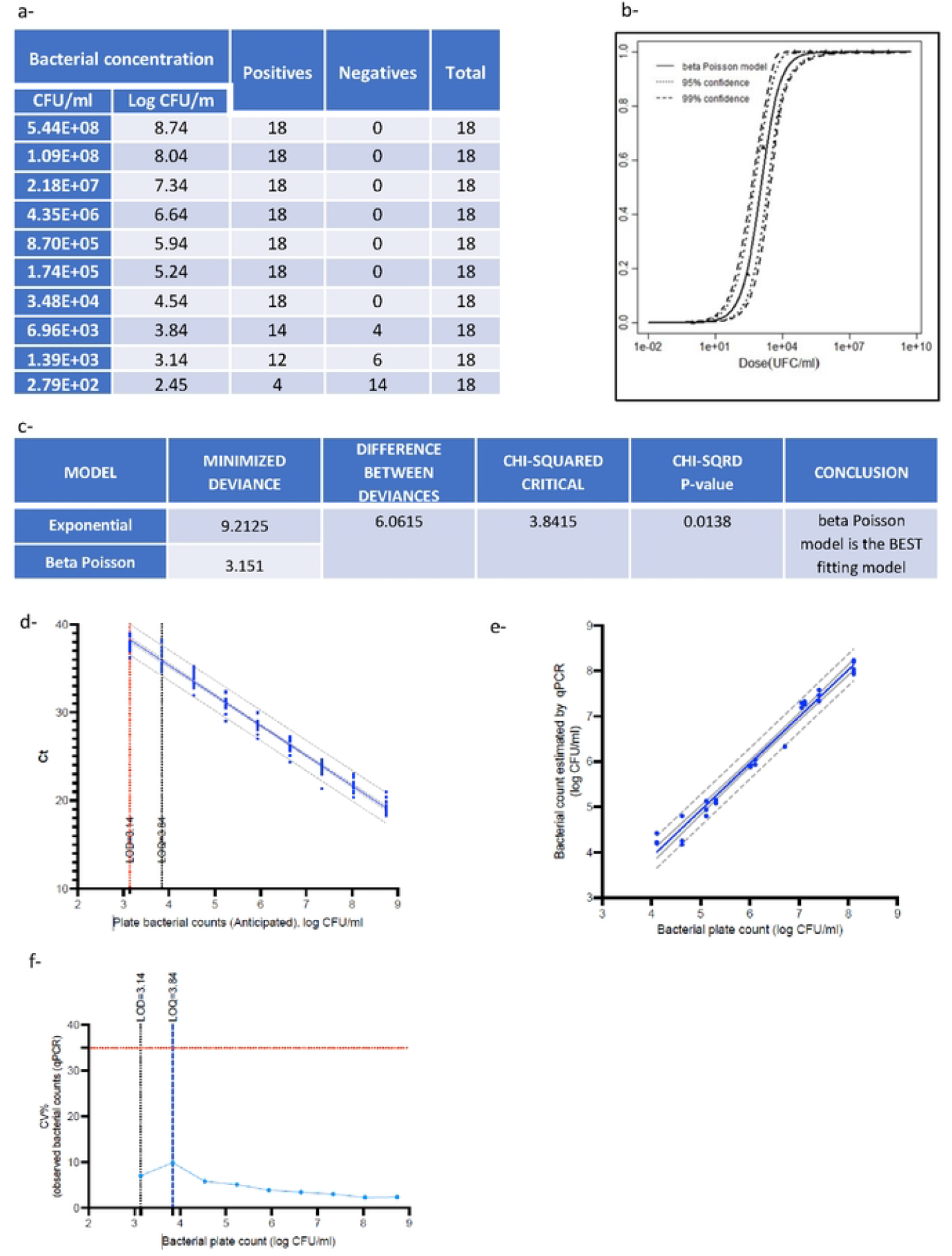
Optimization of qPCR for *Bradyrhizobium diazoefficiens* quantification. (a) Standard curve: Detection rate of 10 serial 5-fold dilutions of bacterial DNA, each tested in six independent qPCR assays with three eplicates per dilution, (b) Detection rate as a function of the bacterial dose (CFU/mL). The solid line represents the Beta-Poisson model, and the dashed lines show the 95% and 99% confidence intervals. Parameters of the Beta-Poisson model (alfa and N50) and LOD estimated from the fitted curve are indicated, (c) Comparison of model fit for LCD estimation using Beta-Poisson and Exponential models.(d) Dynamic range of qPCR and consensus standard curve with 95% confidence and prediction intervals. The limits of detection (LOD) and quantification (LOQ) of the assay are indicated.(e) Linear regression between the viable bacterial estimated by PMA-qPCR and bacterial plate count, (f) LOQ estimation based on the curve of the coefficient of variation (CV% = 100 × SD/mean) as a function of bacterial plate counts (log CFU/mL). The horizontal red dashed line indicates CV = 35%, and the vertical black dashed line marks the LOD and the LOQ defined as the lowest bacterial concentration (log CFU/mL) at which CV falls below 35%.

The data were analyzed using two mathematical models for discrete data. Both methods are referred to as mechanistic, as they assume that the bacterial particles in the sample follow a Poisson distribution around the mean dose. While the exponential model assumes a constant binomial probability that a viable bacterial unit survives in the sample matrix and its genome is detectable by the technique, the beta-Poisson model assumes that this probability varies according to a beta distribution [11]. Finally, the specificity of the qPCR assay was evaluated by testing 20 phylogenetically related strains.

### 2.5 Optimization of PMA-qPCR for Bradyrhizobium diazoefficiens quantification

#### 2.5.1 PMA treatment

PMA (PMA xx, Biotium Inc., USA) was diluted in DEPC water (diethylpyrocarbonate-treated, deionized, filtered water, UltraPure, Invitrogen, USA) to a working concentration of 5 mM and stored at -20°C. PMA treatment was performed according to the manufacturer’s instructions. Briefly, 100 µl of PMA Enhancer (Biotium Inc., USA) was added to each 400 µl sample, followed by a volume of 5 mM PMA according to the desired final concentration (50, 75 or 100 µM). Subsequently, the samples were incubated with mechanical shaking at room temperature for 30 min wrapped in aluminum foil to avoid exposure to light. Then, they were exposed to blue LED light (PhAST Blue, Geniul, Spain) for 15 min to achieve photoactivation of the dye. Finally, the excess of PMA was removed by centrifugation at 8000 xg for 10 min (Centrifuge 5417C, Eppendorf, Germany) and the supernatant was discarded.

#### 2.5.2 Experiment 1 - Minimal concentration of PMA to inhibit non-viable cell signal

Exponential phase cultures of *Bradyrhizobium diazoefficiens* were serially diluted and heat-inactivated (95°C for 10 min). The resulting dilutions (4-9 log CFU/ml) were treated with 50, 75 and 100 µM of PMA. DNA was extracted and analyze by qPCR in triplicate.

#### 2.5.3 Experiment 2 - Assessment of potential interference of PMA on the amplification of different concentrations of viable cells

Two aliquots of three serial dilutions of an exponential phase bacterial suspension (6, 7 and 8 log CFU/ml) were prepared. One aliquot was treated with PMA while the other served as control. Data were analyzed using two-way ANOVA, considering PMA treatment (with or without) as a fixed effect and bacterial concentration as another fixed effect. Post-hoc comparisons were performed using Tukey’s test. Statistical significance was set at p < 0.05.

#### 2.5.4 Experiment 3 - Evaluation of the PMA-qPCR’s accuracy in quantifying viable bacteria in the presence of a high concentration of non-viable cells

A *Bradyrhizobium diazoefficiens* culture in exponential phase of 8.64 log CFU/ml was heat-inactivated (95°C for 10 min) and fractionated into 4 tubes, each containing 900 µl. Subsequently, 100 µl of viable bacteria was added and serial 10-fold dilutions were performed using the tubes containing non-viable bacteria. Each tube was treated with PMA as described in 2.5.1. The aim was to evaluate four concentrations of live bacteria (7.64, 6.64, 5.64 and 4.64 log CFU/ml) in the presence of a high concentration of dead bacteria (8.64 log CFU/ml). Data were analyzed using one-way ANOVA, considering bacterial concentration as fixed effect. Post-hoc comparisons were performed using Tukey’s test. Statistical significance was set at p < 0.05.

#### 2.5.5 Experiment 4 -Determination of PMA-qPCR effectiveness in distinguishing viable from non-viable bacteria within the same concentration range

A *Bradyrhizobium diazoefficiens* culture in exponential phase of 8.64 log CFU/ml was heat-inactivated (95°C for 10 min) and fractionated into 4 tubes, each containing 500 µl. Subsequently, 500 µl of viable bacteria was added and serial 2-fold dilutions were performed using the tubes containing non-viable bacteria. Each tube was treated with PMA as described in 2.5.1. The aim was to evaluate four concentrations of live bacteria (8.43, 8.13, 7.83 and 7.53 log CFU/ml) within the same order of magnitude in the presence of a high concentration of dead bacteria (8.64 log CFU/ml). Data were analyzed using one-way ANOVA, considering bacterial concentration as fixed effect. Post-hoc comparisons were performed using Tukey’s test. Statistical significance was set at p < 0.05.

### 2.6 Experiment 5 - Assessment of PMA-qPCR performance in distinguishing viable from non-viable cells across a serial 2-fold concentration range

A total of 3 assays were conducted, each assay included 3 samples of the commercial product. Each sample had different storage time (0, 12, 24 months post-production) and therefore different concentrations of viable and non-viable bacteria. Since the expected concentrations exceeded the dynamic range of the qPCR, we worked with 1:100 dilutions. Samples were analyzed using both, PMA-qPCR and conventional plate count to estimate the number of live bacteria. Finally, a lineal regression analysis was performed to evaluate the agreement between both methods, with the aim of validating PMA-qPCR as a predictive tool for viable bacterial quantification, potentially replacing plate counting in-process quality control of the product. As a control, samples without PMA were included. For the latter, data were analyzed using one-way ANOVA, considering PMA treatment as fixed effect. Post-hoc comparisons were performed using Tukey’s test. Statistical significance was set at p < 0.05.

## 3. Results

### 3.1 qPCR performance parameters

The efficiency values for all assays were within the acceptable range (90-105%), indicating high-quality qPCR data. The LOD, defined as the lowest concentration of DNA that can be reliably detected, was determined to be 3.14 log CFU/ml using the beta-Poisson model. The LOQ, defined as the lowest detectable concentration within the linear range of the standard curve, was found to be 3.84 log CFU/ml. The dynamic range, defined as the range of concentration over which the standard curve is linear, was determined to be between 8.74 and 3.14 Log CFU/ml (Figure 1). The specificity assay showed that none of the analyzed strains generated a qPCR signal (Data not shown). The intra-assay repeatability, expressed as the standard deviation of the replicates at each point of the standard curve, was less than 0.3 in all cases. Similarly, the inter-assay reproducibility, expressed as the coefficient of variation between runs, was less than 10% (Supplementary material).

### 3.2 Experiment 1 - Minimal concentration of PMA to inhibit non-viable cell signal

Five out of six concentrations of non-viable bacteria were undetectable with the three PMA concentrations analyzed (50, 75 and 100 µM). The 9 log CFU/ml concentration yielded a Ct value of 40 which was interpreted as negative since it was above the limit of detection (Ct=38.17). To optimize resources, the lowest PMA concentration was chosen for subsequent assays. Additionally, it was determined that the maximum allowable concentration of non-viable bacteria in the system was 9 log CFU/ml since a fluorescence signal was observed at that concentration (data not shown).

### 3.3 Experiment 2 - Assessment of potential interference of PMA on the amplification of different concentrations of viable cells

Analysis revealed no statistically significant differences in viable bacteria counts between PMA-treated and untreated samples for any of the concentration tested (6-8 log CFU/ml). These results indicate that PMA did not interfere with the signal from viable bacteria (Figure 2a).

**Figure 2.**
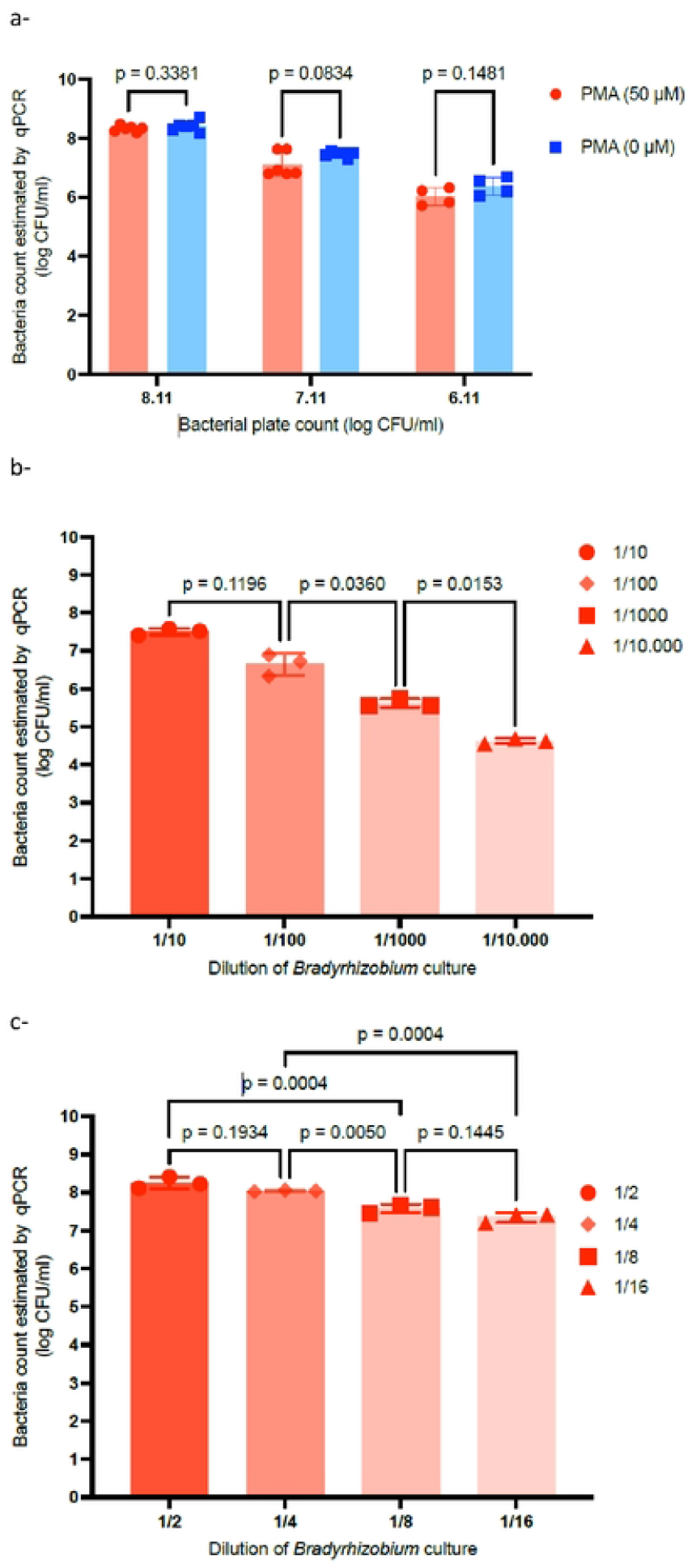
Optimization of PMA-qPCR for quantifying *Bradyrhizobium diazoefficiens*. (a) Experiment 2: Assessment of potential interference of PMA on the amplification of different concentrations of viable cells, two-way ANOVA, Tukey. (b) Experiment 3: Evaluation of the PMA-qPCR’s accuracy in quantifying viable bacteria in the presence of a high concentration of non-viable cells (serial 10-fold dilutions of the bacterial culture; 1/10 = 7.64 log CFU/mL viables in 8,64 log CFU/ml non-viable bacteria), (c) Experiment 4: Assessment of PMA-qPCR performance in distinguishing viable from non-viable cells across a serial 2-fold concentration range (dilution of the culture 1/2 = 8.43 log CFU/mL viables in 8,64 log CFU/ml non-viable bacteria). One-way ANOVA-Tukey.

### 3.4 Experiment 3 - Evaluation of the PMA-qPCR’s accuracy in quantifying viable bacteria in the presence of a high concentration of non-viable cells

PMA-qPCR significantly distinguished among 10-fold serial dilutions (7.64, 6.64, 5.64 and 4.64 log CFU/ml counts), even with high levels of non-viable bacteria (one-way ANOVA; p<0.001) (Figure 2b).

### 3.5 Experiment 4 - Determination of PMA-qPCR effectiveness in distinguishing viable from non-viable bacteria within the same concentration range

PMA-qPCR results were inconsistent when discriminating between 2-fold serial dilutions (8.43, 8.13, 7.83 and 7.53 log CFU/ml). Nevertheless, the technique showed good sensitivity in discriminating 4-fold serial dilutions (one-way ANOVA, Tukey) (Figure 2c).

### 3.6 Experiment 5 - Assessment of PMA-qPCR performance in distinguishing viable from non-viable cells across a serial 2-fold concentration range

Following the linear regression analysis, it was observed that R^2^ was 0.82, indicating that the model explains 82% of the variability of the dependent variable. The regression line, the confidence bands and the 95% prediction bands are illustrated in Figure 3a. As a control, samples without PMA were included. Unlike the PMA-treated samples, they showed no statistically significant differences across the analyzed time points (Figure 3b).

**Figure 3.**
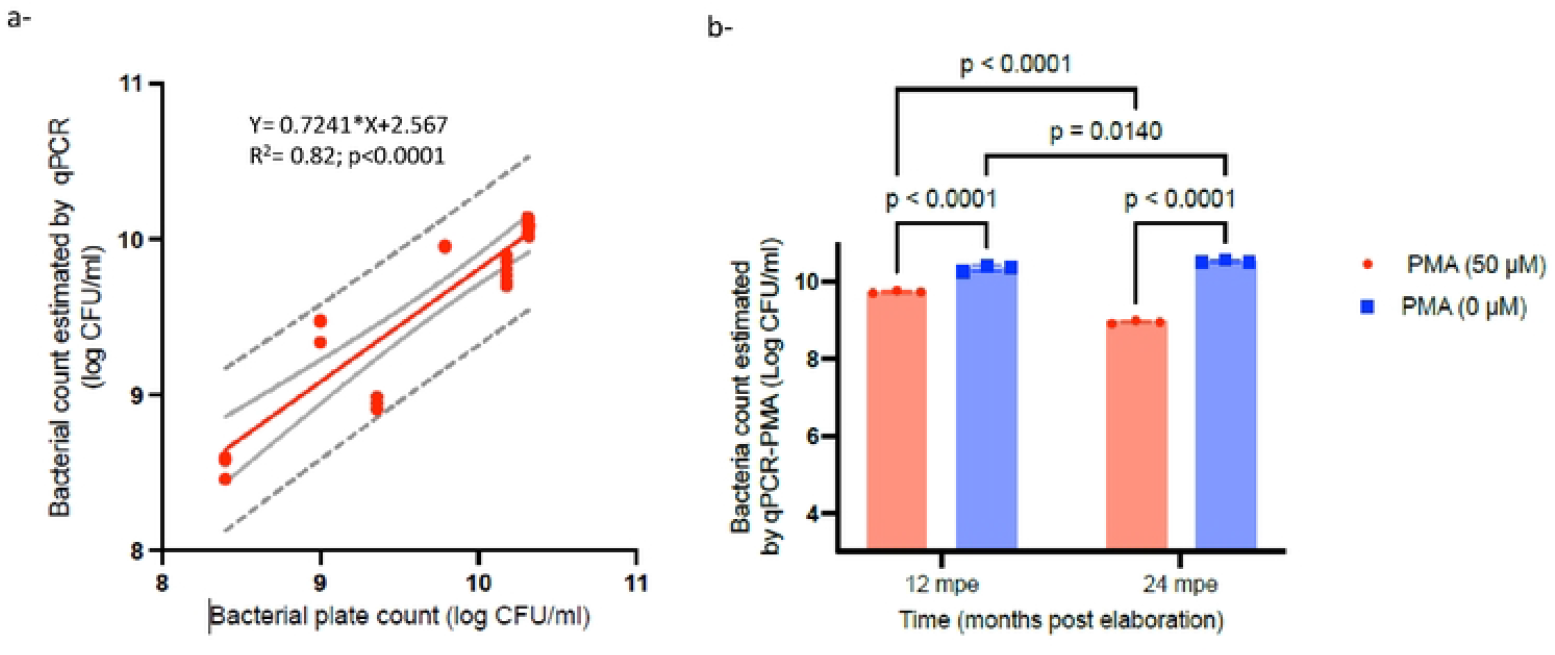
Validation of the methodology using batches of the final product at three different storage times (0-, 12- and 24-months post-manufacturing). (a) Lineal regression between the viable bacteria estimated by PMA-qPCR and the bacterial plate count, (b) Comparison of mean bacterial counts between different storage times (12- and 24-months post elaboration), with and without PMA treatment, repeated measure Anova-Tukey.

## 4. Discussion

Quantification of *Bradyrhizobium* in inoculants currently relies heavily on cultured-based assays, which are time-consuming and labor-intensive. However, accurate quantification is essential to ensure the quality and efficacy of these inoculants. Therefore, the development and validation of rapid, reliable and high-throughput methods, such as the qPCR, are critical for practical agricultural applications.

In this study, we validated a PMA-qPCR assay for the quantification of viable *Bradyrhizobium* as an alternative to conventional culture methods. The performance of the qPCR was demonstrated through the following parameters: an efficiency of approximately 95%, intra-assay repeatability and standard deviation of 0.3, and an intra assay reproducibility with a coefficient of variation less than 10%. These results confirm the reliability and robustness of the qPCR technique for estimating *Bradyrhizobium diazoefficiens* counts.

The LOD and LOQ were also estimated, reflecting the sensitivity of the technique. Although they were slightly higher than those reported for other applications, they are well suited for the intended purpose of this study [12]. A more precise determination of LOD and LOQ could be achieved by performing serial 2-fold dilutions from a 1:1000 dilution onward, following CEFAS recommendations [13]. However, this was beyond the scope of this work, as the primary goal was to estimate the proportion of viable and non-viable cells for batch-to-batch quality control of a product containing viable cell counts in the order of 10 log CFU/ml to 8 log CFU/ml, far beyond the conservative LOD selected for the assay.

Optimizing the concentration of PMA is a crucial step that depends heavily on the intended use of the technique. In our case, samples at time zero were expected to have a viable cell concentration of 10 log CFU/ml. After two years of storage, samples were expected to have a viable cell concentration of 8 log CFU/ml and a non-viable cell concentration of 9.99 log CFU/ml. Thus, the PMA had to effectively inhibit the signal from a high number of non-viable cells in the presence of a high number of viable cells. Since these concentrations exceeded the dynamic range of the qPCR, we worked with 1:100 dilutions, resulting in samples with 6 log CFU/ml of viable cells and 7.99 log CFU/ml of non-viable cells.

To determine the minimum PMA concentration required for complete inhibition of signal from non-viable bacteria, we first tested non-viable cells alone across a range of concentration of PMA concentrations (4-9 log CFU/ml, Experiment 1). Once the minimum PMA concentration was selected, it was evaluated against different concentrations of viable bacteria to ensure it did not interfere with their accurate quantification (Experiment 2). Finally, various concentrations of viable cells were assessed in the presence of a high concentration of non-viable cells (Experiment 3). These assays demonstrated that PMA-qPCR could effectively distinguish among 10-fold serial dilutions (7.64, 6.64, 5.64 and 4.64 log CFU/ml counts), even in the presence of high levels of non-viable cells (8.64 log CFU/ml). To validate the technique’s discrimination threshold a series of twofold dilutions were conducted (Experiment 4). However, while it failed to distinguish between half strength dilutions, it successfully differentiated quarter strength dilutions. This result was expected, as the technique validation had already demonstrated a standard deviation of 0.3 log CFU/ml. Given that the difference between two samples diluted to half-strength is also 0.3 log CFU/ml, the findings are consistent with previous assessments.

As a final assessment (Experiment 5), validation of the method was carried out against the standard plate counting technique, achieving a strong linear correlation with an R^2^ of 0.82. In other words, PMA-qPCR method was able to determine the number of viable cells in only just 5 h, compared to the 120 hours required by conventional plate counting, while maintaining an 82% correlation with the reference method.

To the best of our knowledge, this study is the first to apply PMA-qPCR specifically for quantifying *Bradyrhizobium diazoefficiens* in agricultural inoculants. Pedrolo et al. (2024) evaluated this technique for *Herbaspirillum seropedicae*, an endophytic diazotroph, in pure culture and maize roots, and demonstrate that PMA-qPCR is a powerful approach for quantifying viable and viable but nonculturable cells (VBNC) in inoculants and maize roots. In contrast to the aforementioned study, our results showed no significant differences between plate counts and concentrations estimated from the standard curve. This is likely due to the matrix being specifically formulated to maintain bacterial viability, minimizing the transition to VBNC. The presence of cryoprotectants and appropriate substrates may further prevent VBNC conversion [14].

Another challenge faced by inoculant producers is that if the culture medium becomes contaminated or if inoculants are applied to non-sterile materials, traditional quantification techniques become difficult. This is due to the variable presence of Gram-positive bacteria and fungi, which can interfere with counting accuracy. Some researchers, have proposed improvements to culture media to make them more selective [15,16]. However, none have achieved complete elimination of the accompanying microbiota. In this context, the PMA-qPCR technique stands out for its high specificity.

## 5. Conclusions

This study successfully validated a PMA-qPCR assay for rapid and reliable quantification of viable *Bradyrhizobium diazoefficiens* in inoculants, offering a strong alternative to traditional plate counting. The method demonstrated high efficiency, reproducibility, and selectivity, effectively distinguishing viable cells even in the presence of high non-viable cell concentrations. Validation against plate counting confirmed an 82% correlation, reducing processing time from 120 hours to just 5 hours. These findings highlight PMA-qPCR as a valuable tool for microbial viability assessment in industrial applications.

## ACKNOWLEDGMENTS

This study was funded by private funds from Rizobacter S.A. and public funds from the research project 2019-PE-E7-I120 of the National Institute of Agricultural Technology. We appreciate the participation of Solange Galeano for technical assistance.

## Notes

### Competing Interest Statement

The authors have declared no competing interest.

